# Algorithms for a Commons Cell Atlas

**DOI:** 10.1101/2024.03.23.586413

**Authors:** A. Sina Booeshaghi, Ángel Galvez-Merchán, Lior Pachter

## Abstract

Cell atlas projects curate representative datasets, cell types, and marker genes for tissues across an organism. Despite their ubiquity, atlas projects rely on duplicated and manual effort to curate marker genes and annotate cell types. The size of atlases coupled with a lack of data-compatible tools make reprocessing and analysis of their data near-impossible. To overcome these challenges, we present a collection of data, algorithms, and tools to automate cataloging and analyzing cell types across tissues in an organism, and demonstrate its utility in building a human atlas.

## Introduction

Cell atlas projects such as the Human Cell Atlas (HCA) aim to produce “reference maps” for all cells in the human body. Specifically, the stated goal of the HCA is To create […] reference maps of all human cells […] as a basis for both understanding human health and diagnosing, monitoring, and treating disease”. This aim, shared by various atlas projects such as Azimuth (Hao et al. 2023) and Tabula Sapiens (Tabula Sapiens Consortium* et al. 2022), entails generating a catalog of cell types, states, locations, transitions, and lineages in all cells in the human body using a variety of data sources.

These current atlas projects have multiple drawbacks that limit the scale of data preprocessing and reference map creation. First, data are often preprocessed with different tools introducing unnecessary computational variability (Davis et al. 2018; Z. Zhang et al. 2021; Booeshaghi, Sullivan, and Pachter 2023). Second, quantifications are often limited to the gene-level and do not distinguish between spliced and unspliced molecules (Hjörleifsson et al. 2022). Third, compatible tools for reanalyzing processed data are lacking. Fourth, manual and time-consuming effort is required to annotate cell-types from marker gene lists. Fifth, atlas infrastructure is fixed, limiting the ease with which new cell-type markers and reference transcriptomes can be used to update quantifications.

These issues are apparent in an examination of the HCA. While the HCA provides access to datasets from 427 projects comprising a total of 54.8 million cells (“HCA Data Explorer” 2024a), the data has been processed by a variety of tools, only some of which are maintained and supported by the HCA. For example, while the 10xv2 and 10xv3 data hosted at the HCA portal was processed by Optimus (“Optimus Overview” 2024), datasets such as the human pancreas dataset on the HCA portal (“HCA Data Explorer” 2024b) from (Baron et al. 2016), which is an inDrop single-cell RNA-seq data, was processed with the inDrop workflow (Klein et al. 2015). The CEL-Seq2 pancreatic dataset, also in the HCA portal (“HCA Data Explorer” 2024c) from (Muraro et al. 2016), was processed with a custom BWA workflow. Some datasets, such as (“HCA Data Explorer” 2024d) from (Shrestha et al. 2021), reference the use of multiple different versions of Cell Ranger (v2.1 and v3.1) making reproducibility challenging. The application of these inconsistent preprocessing pipelines extends to much of the HCA data. For example, of the 5,947,388 pancreas cells in the HCA portal, only 51,835 (0.87%) were processed uniformly with the HCA Data Coordination Platform’s (DCP) uniform pipeline. Because the HCA data is not processed uniformly, joint analysis of the data would be very challenging (Tyler, Guccione, and Schadt 2023; Antonsson and Melsted 2024), and the HCA provides neither tools nor recommendations for how to jointly analyze samples (**Supplementary Table 1**). The HCA acknowledges these challenges resulting from the upload of contributor-generated count matrices noting that “[processing] techniques are used at the discretion of the project contributor and vary between projects (“HCA Data Portal Data Matrix Overview” 2024). The HCA also does not provide a curated list of marker genes across cell types by tissue, and lack of standardization of gene annotation references and gene naming complicates comparisons between datasets. Furthermore, most of the HCA quantifications are provided at the gene-level, precluding isoform-level analysis of results (Joglekar et al. 2021; Booeshaghi et al. 2021) or analyses requiring separate quantification of nascent and mature molecules (Gorin, Vastola, and Pachter 2023). Similar problems afflict other single-cell atlas projects such as the Chan Zuckerberg Initiative (CZI) cellxgene atlas (Megill et al. 2021). The HCA and CZI examples, along with the increasing amount of single-cell data that is being generated, illustrates the importance of solving these challenges, specifically of enabling uniform data (re)processing and reference map creation.

In order to overcome these drawbacks in current atlas design, we introduce a set of algorithms, tools, and infrastructure that enables uniform preprocessing with tools that are data-compatible, and with infrastructure that can easily incorporate updated cell-type definitions, references, and which is readily extensible to other modalities. This infrastructure underlies the Human Commons Cell Atlas.

## Results

To facilitate the generation of reference maps from single-cell genomics data, we developed a collection of tools, *mx* and *ec*, that operate on cell by feature (gene/isoform/protein/peak) matrices and marker equivalence class files respectively and work together with uniform preprocessing tools *kallisto, bustools, kb-python*, and *cellatlas (Booeshaghi, Sullivan, and Pachter 2023; Melsted et al. 2021; Melsted, Ntranos, and Pachter 2019; Bray et al. 2016)* (Methods). These tools solve key algorithmic and infrastructure problems in single-cell RNAseq preprocessing, namely automated cell type assignment, marker gene selection, and iterative data reprocessing.

The first step in single-cell data analysis is filtering out low quality cells. This is usually achieved by using a *knee plot*, where the user has to visually find the inflection point that separates good from bad cells. However, this method is manual and subjective, and therefore unsuitable for automated and reproducible data analysis. We solved this problem by implementing ‘‘*mx filter’*, which runs a Gaussian Mixture Model on the 1D histogram of the UMI counts. The tool uses the fact that inflection points in a knee plot correspond to points between peaks in the 1D histogram (**Figure 1a,b,c)**. Therefore, by finding the points of maximum entropy (i.e. maximum uncertainty for the GMM model), *‘mx filter’* can identify the corresponding knee and use it to filter out cells in an automated and efficient way.

**Figure 1:**
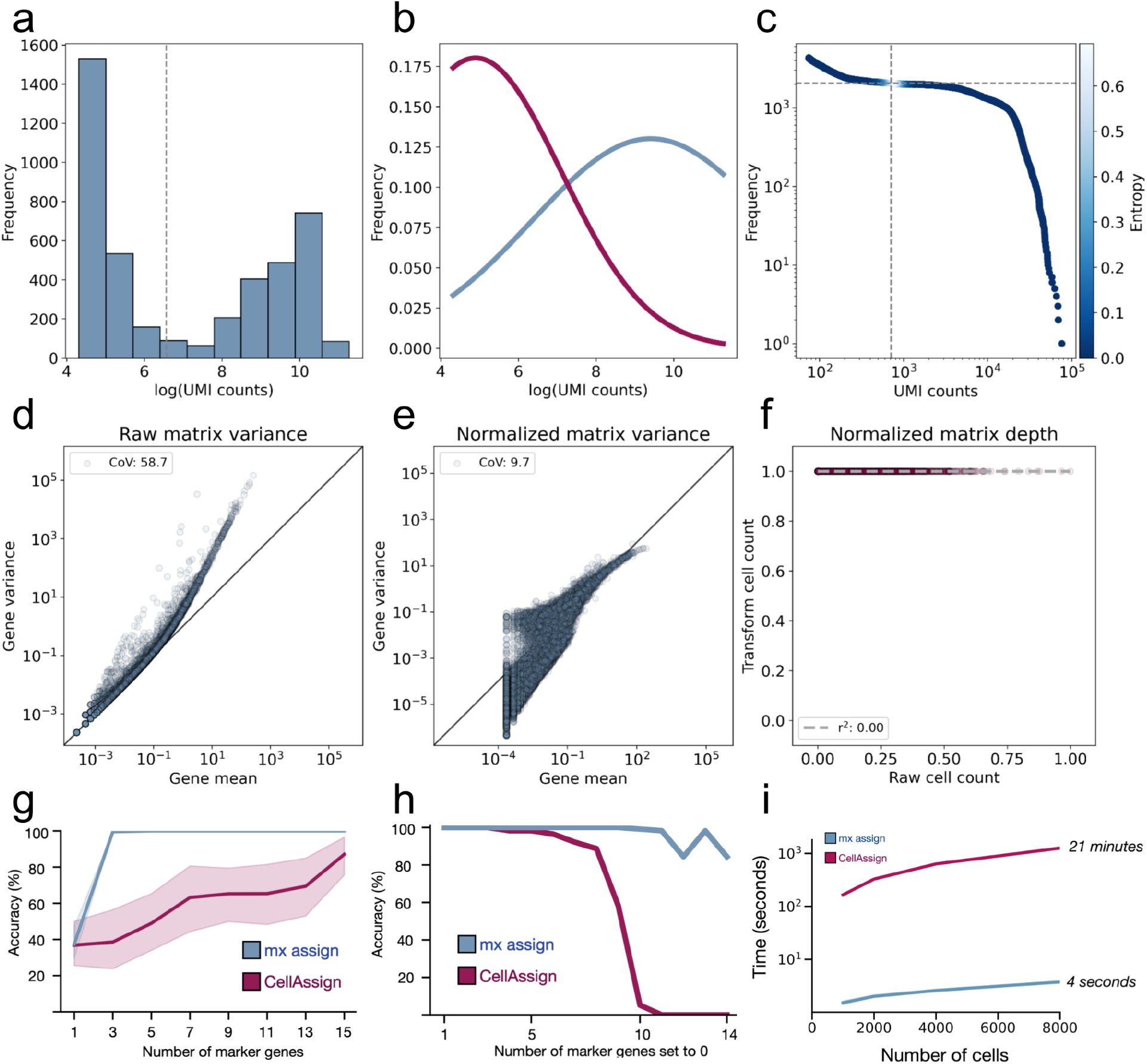
Cell atlas benchmarks. (a) The distribution of UMI counts per cell for cells assayed in colon (GSM3587010). (b) The two Gaussian distributions inferred by the GMM from the UMI counts. The blue line corresponds to Gaussian with a high mean and the red line to the Gaussian with the low mean. (c) The knee plot, where for each cell the UMI counts are plotted on the x-axis and the rank of the cell on the y-axis. The cells are colored by their entropy of assignment to one of the two Gaussian distributions fit by the filtering procedure. The cells are filtered at the point of maximum entropy-colored in white and marked by the horizontal and vertical dotted lines. (d) The mean-variance relationship for genes assayed in cells from the colon (GSM3587010) on the raw matrix. (e) The mean variance relationship on the PFlog1pPF transformed matrix. (f) The cell depth on the transformed vs raw matrix. (g) Accuracy of cell assignment with a varying number of marker genes. Blue is mx assign and red is CellAssign. (h) Accuracy of cell assignment with a varying number of marker genes synthetically set to zero (dropped) in the count matrix. (i) Runtime of cell assignment for a varying number of cells.

After filtering, normalization must be applied in order to remove the confounding technical effects related to the mean-variance relationship of gene expression, as well as those related to variable cell-read depth. We chose the PFlog1pPF normalization (Sina Booeshaghi et al. 2022) as it effectively removes the mean-variance relationship while preserving depth normalization (**Figure 1d,e,f)**.

Next, we aimed to address the problem of manual cell-type assignment. Current methods match differentially expressed genes found on de-novo clustered cells to marker genes from the literature to label cell types. In addition to being time-consuming, this procedure suffers from the double-dipping problem where statistically significant genes are found by testing on groups of cells that are, by construction, different on those sets of genes. Automated cell-type annotation tools such as CellAssign (A. W. Zhang et al. 2019), which operate on a predefined list of marker genes for each cell type, overcome this problem but still suffer from long runtimes and high memory usage. We developed *‘mx assign’* which takes in a single-cell matrix and a marker gene file and performs cell-type assignment using a modified Gaussian Mixture Model. The ‘*mx assign’* algorithm operates on a submatrix of marker genes, like standard algorithms such as CellAssign, but is different in two ways. First, instead of joining multiple batches together to perform assignment, we instead perform assignment on a per matrix basis (we term each matrix an “observation”, Methods). Second, instead of performing assignments on the UMI count matrix, mx assign performs assignments on matrices normalized using ranks (Vargo and Gilbert 2020; Franzén, Gan, and Björkegren 2019; Mei, Li, and Su 2021). This means the distance measurement via Euclidean distance in the GMM is replaced with the Spearman correlation (Methods).

We benchmarked ‘*mx assign’* against CellAssign on simulated data generated using the Splatter R package. We first assessed the accuracy of CellAssign and ‘*mx assign*’ on a varying number of marker genes across different numbers of cell types. We found that ‘*mx assign*’ accurately assigns cells to cell types with as few as three marker genes per cell type while CellAssign performs less accurately on fewer genes (**Figure 1g, Supplementary Figure 1**). Next, we tested the robustness of each algorithm to the mis-specification of marker genes. We simulated the selection of a *bad* marker gene by setting the counts for that gene to 0 for all cells in the sample. We observed that CellAssign loses the ability to correctly assign the cell-type after the misidentification of 10 marker genes, while mx assign remains highly accurate even with a single correct marker and 14 incorrect ones (**Figure 1h, Supplementary Figure 2,3**). Finally, we benchmarked the runtime of each algorithm as a function of the number of cells to be assigned. *mx assign* was 350 times faster than CellAssign at assigning 8,000 cells, demonstrating the efficiency of our algorithm (**Figure 1i**).

Given cell assignments, the next challenge is to perform differential expression at scale for the purpose of cell-type marker identification. Finding cell-type marker genes requires both performing differential expression and selecting marker genes, tasks which are often not clearly delineated, despite being distinct (Sina Booeshaghi et al. 2022). We use a t-test (Soneson and Robinson 2018) on the matrix of all genes to find differentially expressed genes on a different set of genes than those used for cell-type assignment. By performing differential expression on a different set of genes, we avoid the double-dipping problem that current DE methods face.

We then select cell-type marker genes from the list of differentially expressed genes. The identification of marker genes requires selecting genes that are differentially expressed between cell types and highly expressed within cell types. Often lists of differentially expressed genes are reported as marker genes despite marker genes having stricter requirements (**Supplementary Note**). To find marker genes, we implement a gene selection strategy, ‘*ec mark*’, that filters on both the p-value and log-fold change in addition to selecting the most highly expressed genes within the cell type by average gene expression rank.

With a fully-automated method for assigning cell types and generating cell-type marker genes we set out to build a workflow and associated infrastructure that can easily incorporate new data and marker gene lists. Our suite of tools, including *mx, ec*, and *kallisto bustools*, include other key steps in scRNA-seq processing, such as quality control *‘mx inspect’* as well as other more general matrix manipulation procedures whose utility extends beyond single cell matrices. With all these tools in hand, we set to design a workflow and associated infrastructure for the reproducible generation of a single cell atlas, which we named the Commons Cell Atlas (CCA). The workflow, illustrated in **Figure 2**, consists of a series of ‘*mx*’ and ‘*ec*’ commands that can efficiently and modularly process count matrices. The final output of the workflow; the filtered, normalized and assigned matrix, constitutes the unit of the Commons Cell Atlas, which can be constantly modified by the automated and continuous running of the tools. The output also enables the creation of atlas-level matrices that can be used for cross-sample/organ/atlas analysis (**Supplementary Figure 4**). The infrastructure is hosted on GitHub, making any Commons Cell Atlas easily accessible, as well as highly collaborative. We envision the creation of Commons Cell Atlases for many organisms, as well as for many purposes, each of which will grow and evolve as we learn more about the system of study.

**Figure 2:**
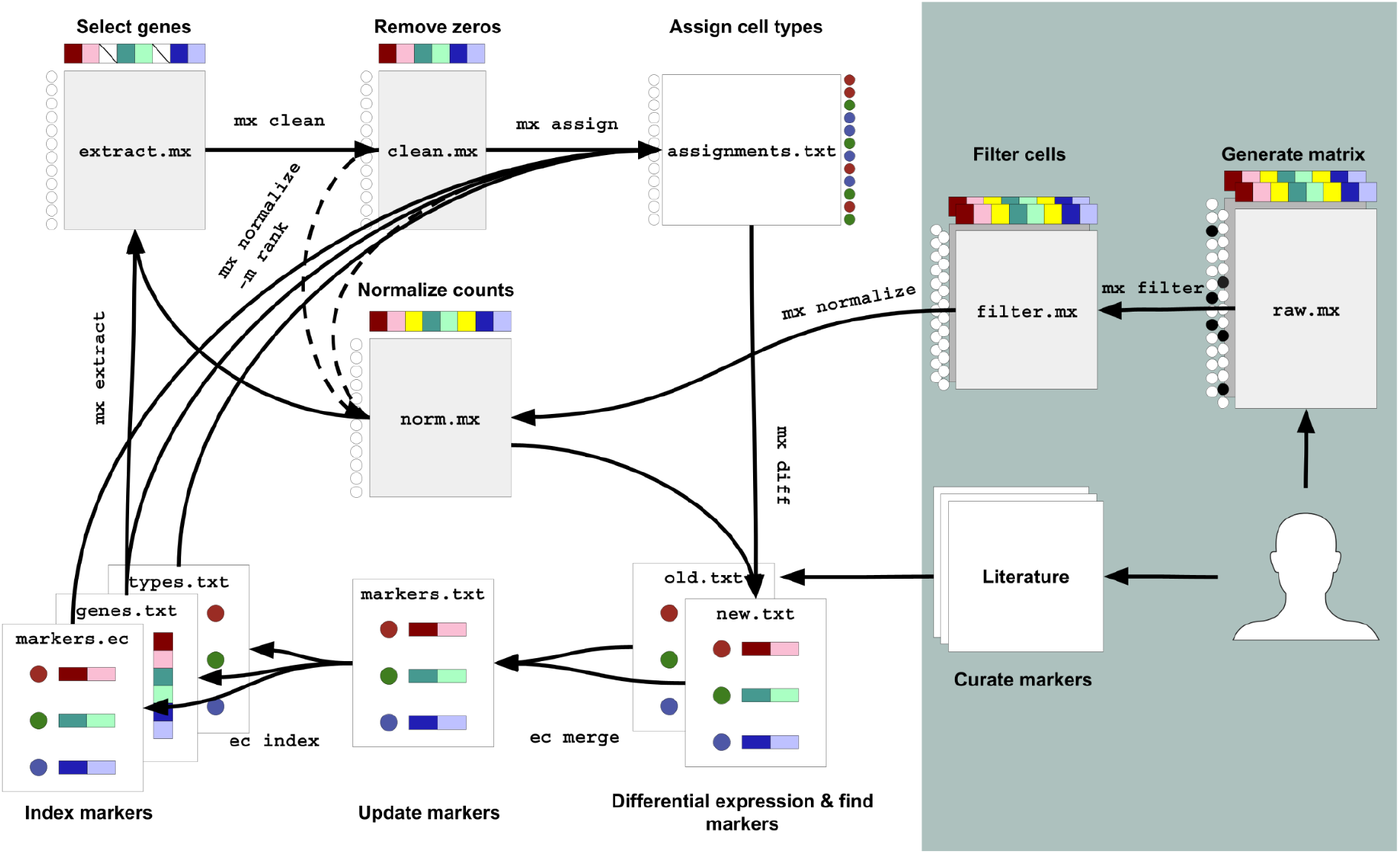
Cell atlas workflow and infrastructure. The cellatlas workflow and infrastructure facilitates matrix normalization and celltype assignment based on marker genes derived from literature. A user supplies a raw count matrix and points to a set of marker genes curated from literature. The user runs the *mx filter* command to filter out cells with low counts by the knee plot method. The user then runs *mx normalize* to perform depth normalization and variance stabilization to the matrix with log1pPF. In parallel, the user runs multiple *ec* commands to take a list of curated markers and produce an indexed marker file to use for cell assignment. After generating the indexed marker file, the user runs *mx extract* and *mx clean* to produce a submatrix that contains only cells with non-zero counts on the genes contained in the marker gene index. Cell assignment is then performed with *mx assign*. Differentially expressed markers are found with *mx diff* and old markers are updated with newly found markers using *ec*. The atlas infrastructure is iterative and flexible. With new marker genes, cell-type assignments can be updated and new markers can be found.

### Open Data

The concept of open data has been central to open science single-cell atlas efforts such as the Human Cell Atlas project (Regev et al. 2018). While there is a general understanding of the significance of open data, e.g. improving transparency, accelerating research, and facilitating discovery of datasets (Vayena and Gasser 2016; Knoppers et al. 2023), there is no precise definition of the term (Bahlai et al. 2019). In the context of the CCA, we have been intentional about open data, specifically engineering the atlas to include open access to underlying data (both raw data and uniformly processed count matrices), key analyses, including lists of marker genes, and accompanying open source tools for exploring the data. The CCA contains 20.74Tb of raw sequencing data (**Figure 3**) in addition to marker genes for 31 tissues derived from previous studies; all of these are available on the public GitHub repository and Zenodo. The open data underlying the CCA can be distributed in this way thanks to authors of prior studies who have made their data publicly accessible. Thanks to this layering of open data, the CCA can be adapted to include other multimodal datasets in the future, such as those generated with 10x Multiome, or datasets from different species into the atlas for analysis. Genes from publicly available tables of differentially expressed genes can also be incorporated into existing CCA cell-type annotations.

**Figure 3:**
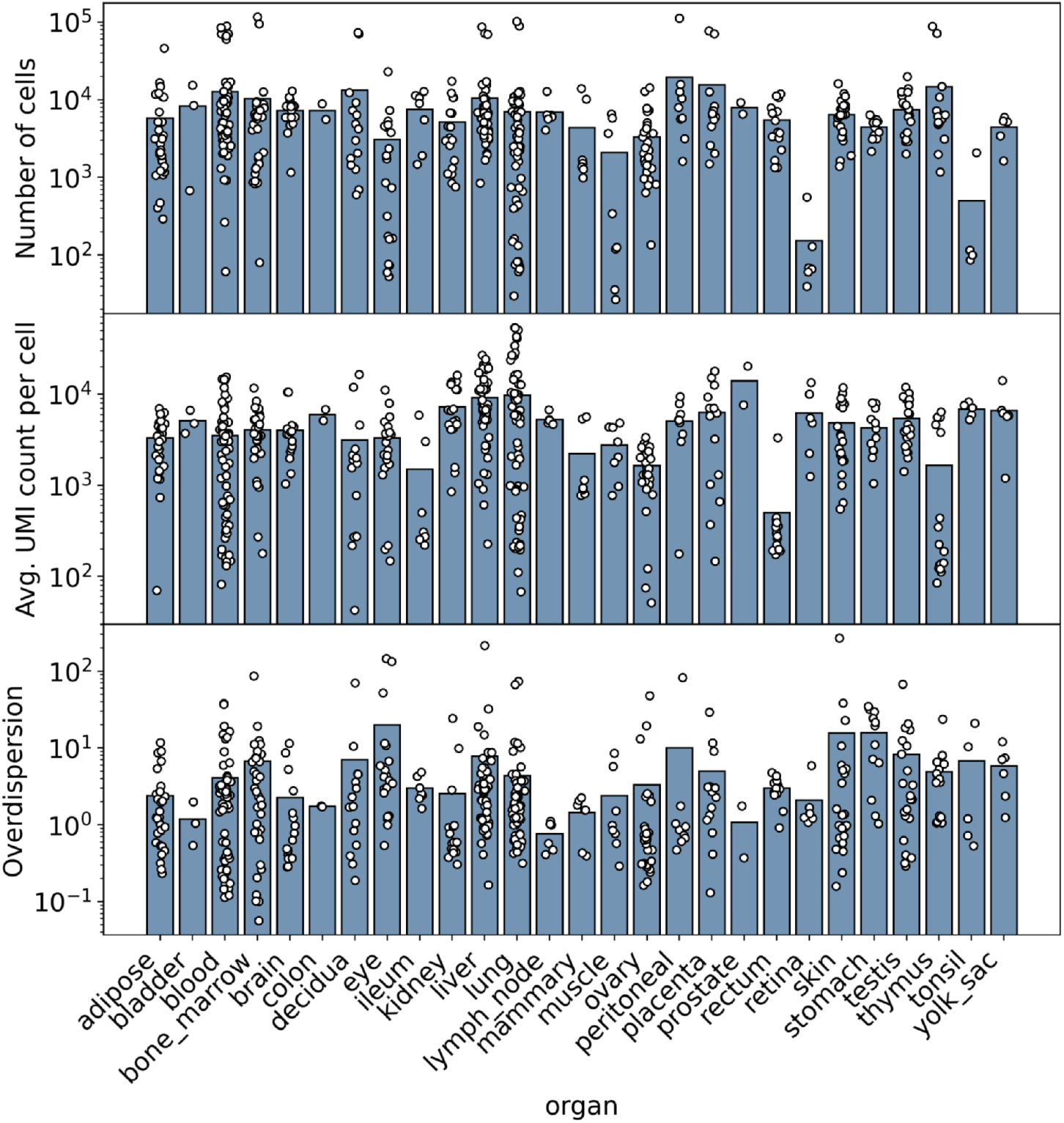
Atlas-level QC metrics generated with the CCA instrstructure. Multiple matrices were preprocessed for each tissue and QC metrics were generated using the *mx inspect* command. Each point corresponds to a dataset (observation, see Methods). The first row shows the number of cells (after filtering) for each matrix. The second row shows the average UMI counts per cell for all cells in each matrix. The third row shows the overdispersion parameter as computed by fitting a quadratic to the mean and variance of expression for each gene using all genes. In all three plots the bar represents the average.

We have strived to impose as few limitations to access to the CCA data as technically possible. FASTQ files can be large and so we point to them via their accessions on publicly available databases. Matrices generated from preprocessing tasks can also be large but we have made them available for users to explore directly via GitHub. We therefore believe our efforts are well characterized by the “Open Data” *Point for Consideration* proposed by the HCA (Regev et al. 2017).

## Conclusion

We’ve developed a series of validated tools and associated infrastructure to routinely process single-cell genomics data in a fast and iterative manner with validated methods. Our tools allow users to go from a collection of FASTQ files to annotated count matrices-a crucial first set of steps that precede all downstream analysis tasks. Importantly, we perform our preprocessing steps in a batch-by-batch manner so as to avoid the introduction of technical variability when combining matrices across batches (Tyler, Guccione, and Schadt 2023). Our tools and infrastructure support the generation of a Human Commons Cell atlas and provide a framework for generating Commons Cell atlases of other organisms and other modalities.

## Discussion and limitations

The notion of a “single-cell atlas” has, over the past decade, gained a specific meaning within the field of genomics. It is widely understood that a single-cell atlas constitutes a collection of single-cell genomics datasets, along with defined clusters of cells that are labeled as cell types according to marker genes. The number of cells comprising an atlas has been used as a measure of its value (“‘tabula Sapiens’ Multi-Organ Cell Atlas Surprising Biologists” 2022). This model of an atlas project as a fixed representation of cells is incomplete without compatible algorithms and tools for data processing and analysis that can facilitate exploration of the atlas. In addition, this model fails to capture the dynamic nature of cellular processes-their life, death, and state-transitions. Additional limitations are imposed on cell atlases by the technologies currently used to collect data. For example, the average depth per cell is widely reported to be ∼10k UMIs per cell, however our extensive data processing across many organs demonstrates the number to be closer to ∼5k per cell (**Figure 3**). Lower cell depth coupled with discordant matrix-level operations impact the types of marker genes that are identified. The CCA framework addresses some of these shortcomings by providing transparent infrastructure to assess the content of an atlas, providing tools for exploration of an atlas, and most importantly a framework for updating atlases with addition of new data.

Another challenge that the CCA addresses is the lack of a standardized cell-type nomenclature and cell-type definitions. The lack of standardization makes combining marker genes derived across various resources challenging since marker-cell-type labels are often inconsistent. While we do not tackle the problem of standardizing cell type or marker gene nomenclature, the efficiency of the CCA and its reliance on prior literature to perform cell type assignment offers an automated solution for incorporating domain expertise via the published literature into an atlas. However, a consequence of this CCA approach is that rare celltype discovery may be challenging. We envision that reanalysis efforts, made possible by the CCA infrastructure, will help mitigate this challenge and will enable various strategies for cell-type identification and cataloging (Zeng 2022; Domcke and Shendure 2023).

The dependence of the CCA on cell types as fundamental units acknowledges that groups of cells undergo dynamic life, death, and transitory processes (Zeng 2022), and it is crucial that atlas projects have a way to account for this dynamism. The CCA does not in and of itself address all the challenges that properly accounting for diversity in cell state and type require, but we posit that its flexibility offers a useful starting point.

One potential drawback of the CCA is that it relies heavily on marker genes for cell type assignment. Several methods have been developed to identify marker genes, and the approaches proposed have grappled with the tradeoff between specificity of markers, and the improvements in sensitivity of assigning cells that can be obtained by increasing numbers of combinatorially defined markers (Efroni et al. 2015; Delaney et al. 2019). Also, while marker genes may be useful to identify distinct cell types, they fail to discriminate and characterize cell types present in multiple developmental stages, such as nematocysts in jellyfish (Chari et al. 2021) Moreover, there is a fundamental tension between definitions of markers that rely on relative expression, versus absolute thresholds for markers (**Supplementary Note**). Specifically, there is no definition for marker genes that simultaneously satisfies three desirable properties: scaling, richness, and consistency. This situation is similar to clustering, where there are similar fundamental tradeoffs and an impossibility of satisfying these three desired properties (Kleinberg 2002). Importantly, these theorems do not negate the utility of clustering or of using marker genes in exploring and analyzing single-cell transcriptomics data. Rather, they highlight the need for choosing what properties are important. In other words, standardized and uniform construction of atlases across tissues and organisms requires agreement on the goals of methods prior to the choice of tools. The uniformly processed dataset of the CCA should facilitate exploration of these questions in a controlled setting.

While we have presented a cell atlas for a single modality across 27 tissues, a cataloging of cellular processes requires a more complete atlas that takes into account multiple cellular measurements such as their spatial distribution, chromatin accessibility, and many other measured modalities in addition to higher quality datasets that span all organs in the human body. The infrastructure we present is extensible, by virtue of being fast and reproducible, enabling researchers to build on it. In principle the CCA infrastructure can be extended to catalog other modalities, but we have not tackled these challenges yet. The seqspec standard (Sina Booeshaghi, Chen, and Pachter 2023), which can help in automating preprocessing of different assays, can in principle help with this goal. Despite these challenges, the tools we have developed are all transparent, facilitate reproducible research, are documented making them usable, and have been evaluated in rigorous benchmarking and far surpass the capabilities of current atlas efforts. They should therefore be readily adaptable to other single-cell genomics modalities.

We expect that our CCA standard will be of value in other atlas projects, and that the modular nature of the CCA infrastructure makes possible the easy adoption of some of our tools for other projects. All methods and tools used to build the CCA project are licensed under the BSD-2 license making them freely usable both in academia and industry. We anticipate they will be useful for consortium-level atlas building projects such as those being undertaken by the BICCN (BRAIN Initiative Cell Census Network (BICCN) 2021), BICAN (Kwon 2023), HUBMAP (Jain et al. 2023), IGVF (IGVF Consortium 2023), HCA (Regev et al. 2017), and Tabula Sapiens (Tabula Sapiens Consortium* et al. 2022).

## Methods

### Data simulation

We simulated scRNA-seq data using the R package Splatter with default parameters, 10,000 genes, a DE probability of 0.2 and an equal probability for each of the simulated groups. We performed simulations varying the number of cells (1000, 2000, 4000 and 8000) and the number of groups (2, 4, 6 and 8).

### Assignment benchmarking

We benchmarked CellAssign and mx assign using the simulated data described above. We followed the same approach as (A. W. Zhang et al. 2019) to select marker genes. Both CellAssign and mx assign were run with default parameters for all the simulated datasets varying the number of marker genes used for the assignment.

### Accuracy measures: marker genes per cluster

We calculated the accuracy by comparing the assignment labels of each method to the group ground truth from the Splatter output. We varied the number of marker genes per cell type from 1 to 14 and compared the assignment of cells per cell type between CellAssign and mx.

### Accuracy measures: marker misspecification

In order to assess the ability for each tool to assign cell types with misspecified markers, we randomly chose sets of marker genes and set their expression to 0. We varied the number of markers where the expression of the gene was set to zero (from 15 to 1) in random order and measured the accuracy of the assignment.

### Runtime

For the runtime benchmark, we ran mx assign and CellAssign on the simulated dataset containing 8 groups and 1,000, 2,000, 4,000 or 8,000 cells using 15 markers per group. The runtime was calculated using the results for 3 independent runs for each condition. Benchmarks were performed on an Intel(R) Xeon(R) CPU E5-2697 v2 @ 2.70GHz 12 cores.

### cellatlas

The tools in the CCA infrastructure can all be run with the *cellatlas* command. cellatlas combines preprocessing along with the two processing tools *mx* and *ec* to generate single-cell assignments from FASTQ files. The cellatlas build command produces a list of kb_python commands, which in turn utilizes kallisto and bustools (Bray et al. 2016; Melsted et al. 2021; Sullivan et al. 2023) to preprocess FASTQ files into count matrices. The code is open source and freely available on GitHub: https://github.com/cellatlas/cellatlas.

### mx

mx is a command-line tool written in Python that manipulates Matrix Market exchange (Boisvert, Boisvert, and Remington 1996) matrices. mx can perform key steps in scRNA-seq data processing, as well as more general matrix manipulation steps that can be useful for a variety of applications. The *mx* tool consists of nine subcommands:

1. assign - Run assignment algorithm
2. clean - Drop rows/cols with all zeros
3. convert - Convert matrix
4. diff - Differential Expression
5. extract - Extract submatrix of genes
6. filter - Filter cells from matrix
7. inspect - Inspect matrix
8. normalize - Normalize matrix
9. split-Split matrix by assignments

### ec

ec is a command-line tool written in Python that manipulates ec formatted files. The format of an *ec* file is an (optional) header that begins with a ‘#’, followed by rows of two columns that are tab-delimited. The first column is the “group” or “cell-type”-a string identifying the name of the group and the second column is a list of “targets” or “gene/isoform names or ids” that are separated with a single comma. The *ec* tool consists of five subcommands:

1. index - Index markers.txt file into groups, targets, and ec matrix files
2. mark - Created markers.txt from deg.txt file
3. merge - Merge two markers.txt files (union or intersection)
4. verify - Check whether the provided groups, targets, ec matrix and markers.txt files are correct and compatible with each other
5. filter - Filter out bad genes from markers.txt file

### mx assign

The ‘mx assign’ program assigns cells to cell-types according to a provided marker gene list. The code of ‘mx assign’ is based on the Gaussian Mixture Model (GMM) implementation of the Python package *‘sklearn’*. The input to ‘mx assign’ consists of a table of marker genes and their associated cell types, along with a count matrix. The first step in ‘mx assign’ is to subset the matrix to the marker genes contained in the markers file. Then rows and columns in the matrix with zero UMI counts are removed, and the remaining elements of the matrix are converted into ranks. Specifically, a gene in a cell is assigned a rank based on the UMI count vector for that gene, with the cell with the highest expression for that gene receiving the highest number. Finally, the GMM algorithm as implemented in *sklearn* is applied to the matrix of ranks to assign cells to cell types.

In a standard GMM, the expectation step uses the Mahalanobis distance measure, which modifies Euclidean distance by taking into account the covariance matrix, to compute the probability that each cell belongs to each cluster. Clusters are modeled with Gaussian distributions. When using rank data, Euclidean distance is proportional to the Spearman correlation, and thus the Euclidean distance between a cell as represented by the previously computed ranks for all genes, and the centroid of a cluster as represented by ranks, is proportional to the Spearman correlation between the cell and the cluster centroid. Therefore the procedure of computing the ranks on the counts prior to running the standard GMM effectively modifies assignment to be based on a nonparametric measure (Spearman correlation). This can be seen from observing that if the rank vectors associated to the cell and the centroid are *r(X*_*i*_) and *r(Y*_*i*_) respectively, and *n* is the number of marker genes, then the Spearman correlation *r*_*s*_ is given by:

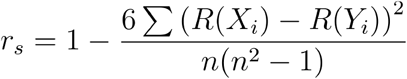

### Atlas infrastructure

Datasets are organized into “runs” and “observation” units. Each run is composed of FASTQ files that correspond to a single sequencing run (demultiplexed on sample index). Observations are sets of runs for which a single count matrix is generated. Oftentimes an observation consists of data from only one run. An observation is associated with various metadata, for example the GEO metadata object returned by *ffq* (Gálvez-Merchán et al. 2022) as well as the tissue of origin and species. The CCA infrastructure is composed of a few key steps

1. Download FASTQs and metadata (*ffq)*
2. Preprocess FASTQ files into count matrices with transcriptomic references (*kb-python)*
3. Generate marker genes from count matrices and marker gene references (*mx & ec)*

A set of software tools facilitate each part of the infrastructure (highlighted in the parenthesis).

Preprocessing is performed with kb-python and uses a species-specific transcriptomic reference to generate a count matrix. mx is used to filter, normalize, and manipulate the count matrix to prepare it for cell-type assignment. ec is used to manipulate the marker gene file for use with mx assign to perform cell-type assignment.

Cell-type assignment is performed in an observation-by-observation manner. First, cell barcodes are filtered out from the count matrix using mx filter (which implements the knee filter). The matrix is then normalized with mx norm. In parallel, ec index is used to prepare the marker gene list for cell-type assignment. The indexed-marker gene file is used to subset the normalized count matrix to only the genes contained in the marker gene file. Resultant cells that have zero expression of all marker genes are cleaned up and mx assign is run. Assignments are reported alongside the Shannon entropy of cell-assignment to all Gaussians. Using the new assignments, mx diff is then run to perform differential expression across all of the genes. By performing cell assignment on a subset of the genes and performing differential expression using the remainder, we aim to avoid the double-dipping problem whereby genes are called significant in DE due to their importance in clustering. Lastly, ec mark is used on the differential expression results to define a new set of marker genes that can be used to update the previous.

### GitHub Usage

We use GitHub as a centralized location to store observation metadata; observations are separated into their tissue of origin. Metrics for each observation are also stored on GitHub. Due to the size limitations imposed by GitHub, storing matrices or FASTQ files on GitHub is not feasible. We therefore store pointers to datasets via their *ffq* metadata. While not implemented in this work, GitHub actions can be used to automate the cell atlas infrastructure. This would make it straightforward to add datasets and automatically get back assignments and marker gene lists.

## Data and code availability

The code and data needed to reprocess the results of this manuscript can be found here https://github.com/pachterlab/BGP_2024/. The CCA atlas can be found here https://github.com/cellatlas/human/. A summary of the datasets in the CCA atlas can be found here https://cellatlas.github.io/human/.

## Author contributions

The CCA atlas concept emerged from an initiative by ASB to uniformly preprocess the datasets in (Svensson, da Veiga Beltrame, and Pachter 2020). ÁGM conceived of the idea of examining the OAS1 isoforms at single-cell resolution across human tissues after the publication of (Zhou et al. 2021). ASB conceived the CCA structure and associated mx and ec toolkit with feedback from LP. ÁGM pre-processed the CCA datasets, and ASB and ÁGM wrote mx and ec. ÁGM and ASB developed the CCA quality control. ÁGM led the OAS1 analysis, with help from ASB and LP. ASB developed the ‘mx assign’ cell assignment approach, and ÁGM and ASB benchmarked it. ASB drafted the initial version of the manuscript, which was edited and reviewed by all authors.

## Acknowledgments

The authors acknowledge the Howard Hughes Medical Institute for funding A.S.B. through the Hanna H. Gray Fellows program. Thanks to the Caltech Bioinformatics Resource Center for assisting with pre-processing the data.

## Competing Interests

None.

## Notes

https://github.com/cellatlas/human/

https://github.com/pachterlab/BGP_2024/

### Competing Interest Statement

The authors have declared no competing interest.

