## Supplementary Note for "Algorithms for a Commons Cell Atlas"

March 23, 2024

In single-cell RNA-seq, cell-type assignment [1] is the task of associating a collection of cells with cell types that are defined in terms of marker genes [2]. Therefore, the assignment depends on identifying, for each cell, the markers it expresses. Formally, *markers* for cell types are encoded as an $n \times k$ binary matrix $M$ where entry $M_{ij}$ specifies whether gene $i$ is a marker for cell type $j$. A *gene expression matrix* is an $n \times N$ matrix $E$ where $E_{ij}$ is a real positive number specifying, in some units, the abundance of gene $i$ in cell $j$ for $n$ genes and $N$ cells. In other words, the markers for a cell type $j$ correspond to subsets of $2^{[n]}$ corresponding to the genes that are present and the expression of each cell is a vector in $\mathbb{R}^n$.

A *cell marker function* is a function that reports for each gene expression vector corresponding to a cell, its markers genes, i.e. a function $f : \mathbb{R}^n \to 2^{[n]}$. For a given marker matrix $M$ and an expression matrix $E$, such a function can be used to identify the markers expressed in each cell, and therefore allow for the association of the cell with one of the cell types. In analogy with the axioms for clustering suggested in [3], we posit that a cell marker function should satisfy:

1. $f(\alpha c) = f(c)$ for a gene expression vector $c$ and every real positive scaling factor $\alpha$. This is called *scale invariance.*

2. For all $A \subseteq 2^{[n]}$, there exists a gene expression vector $c$ such that $f(c) = A$. This is called *richness.*

3. If $f(c) = A \subseteq 2^{[n]}$ for some gene expression vector $c$, and $c'$ is another gene expression vector with $c'(i) \geq c(i)$ when $i \in A$ and $c'(i) \leq c(i)$ when $i \notin A$, then $f(c') = f(c) = A$. This is called *consistency*

Biologically and technically, the scale-invariance property reflects the fact that gene expression matrices are estimates of *relative* abundances of genes. Richness is the mathematical requirement that cell marker functions are surjective. Biologically, this means that in principle, any combination of genes constitutes a set of markers for some kind of cell. Consistency reflects an essential property of markers: if a cell expresses a marker, then if the amount of

*These authors contributed equally
†To whom correspondence should be addressed.

that marker gene increases, the marker gene should still be considered a marker. Similarly, if a gene is not a marker gene for a cell then if the gene is expressed less it should not switch to being a marker gene.

In analogy with the impossibility result for clustering from [3]:

**Theorem 1.** *Let $n \geq 2$. No cell marker function $f : \mathbb{R}^n \to 2^{[n]}$ can simultaneously satisfy scale-invariance, richness and consistency.*

*Proof*: By richness, there must exist a gene expression vector $c$ such that $f(c) = \{1, 2, \dots, n\}$. Since $n \geq 2$, there must exist another gene expression vector $c'$ such that $f(c') \neq f(c)$. Let $\alpha = \frac{max_i c(i)}{min_i c'(i)}$ and let $c''(i) = \alpha c'(i)$. Note that $c''(i) \geq c(i)$ for all $i$, and therefore by consistency $f(c'') = f(c)$. By scale-invariance $f(c'') = f(c')$. Since $f(c)$ and $f(c')$ are both equal to $f(c'')$, it follows that $f(c) = f(c')$, which is a contradiction. $\square$

This proof mimics the proof of Kleinberg's theorem by [4]. It is the formalization of the incompatibility of the *relative* changes facilitated by scale-invariance and the *absolute* differences allowed by consistency if richness is to be required. The theorem does not mean that there is no meaningful approach to defining marker genes. Rather, it highlights choices that must be considered when deciding on a method to select marker genes.

[3] Jon Kleinberg. An impossibility theorem for clustering. *Advances in neural information processing systems*, 15, 2002.

[4] Margareta Ackerman. *Towards Theoretical Foundations of Clustering.* PhD thesis, University of Waterloo, 2012.
