## Supplementary Information for "Algorithms for a Commons Cell Atlas"

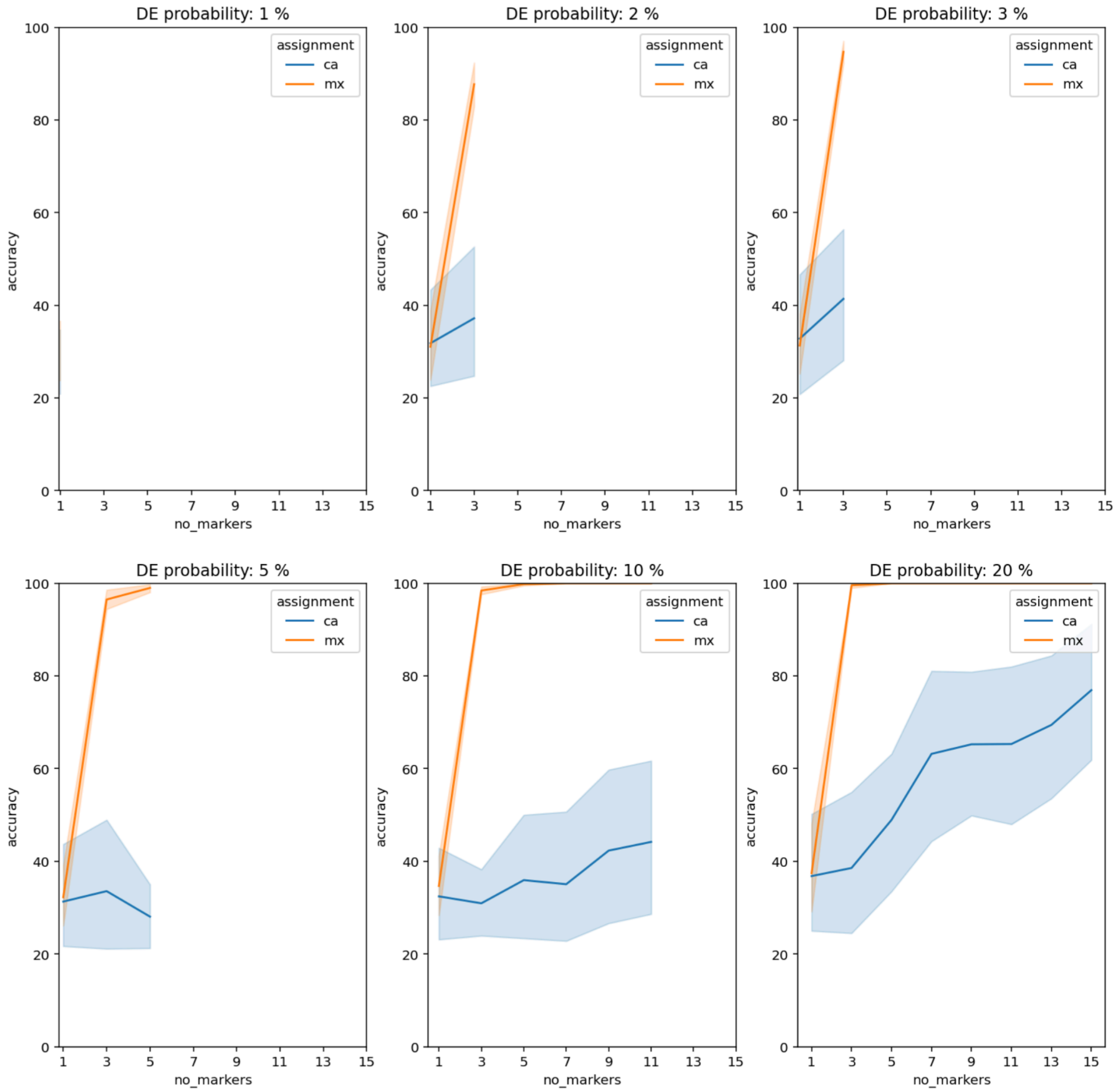


**Supplementary Figure 1:** CellAssign and *mx* were used to assign cell-types on Splatter simulated data across different DE probability values. For each DE probability, we performed simulations with different numbers of cells (1000, 2000, 4000 and 8000) and different number of cell types (2, 4, 6 and 8). CellAssign and *mx* were run using 1 to 15 marker genes, and the assignment accuracy was measured. Marker genes were selected using the same strategy as in CellAssign's original publication (Zhang et al. 2019), genes that were in the top fifth percentile of log-fold change and the top tenth percentile of mean expression. As a consequence of this marker gene selection method, less than 15 markers could be selected for low DE probabilities.

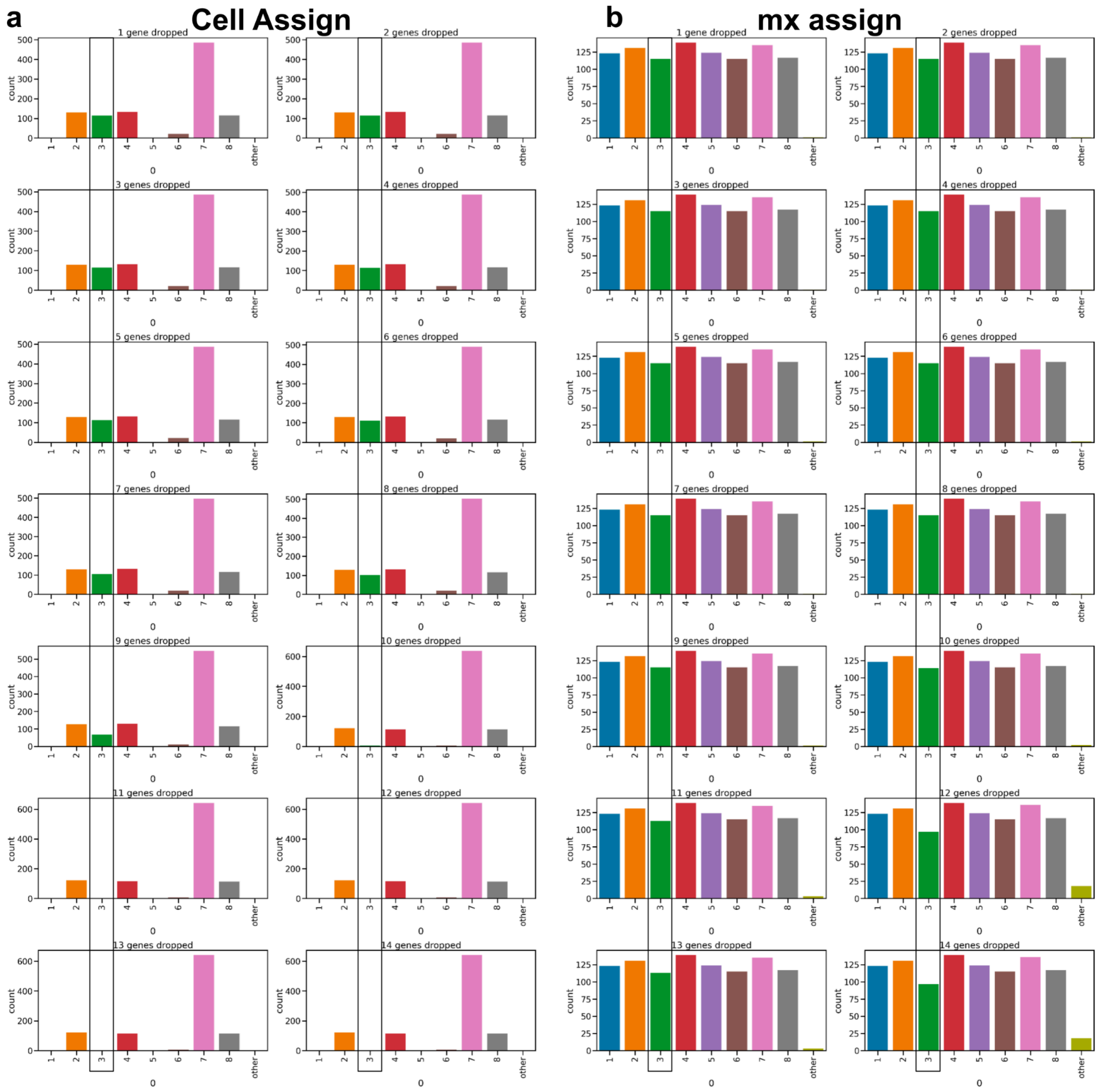


**Supplementary Figure 2:** The number of cells assigned to each cell type by CellAssign (first column) and *mx* (second column). The expression of the marker genes for cell-type 3 was set to 0 sequentially in random order, and the number of cells assigned to each group was measured. An extra cell-type with no marker genes (*Other*) was included in the assignment with the goal of capturing cells that couldn't be assigned to any of the defined cell-types.

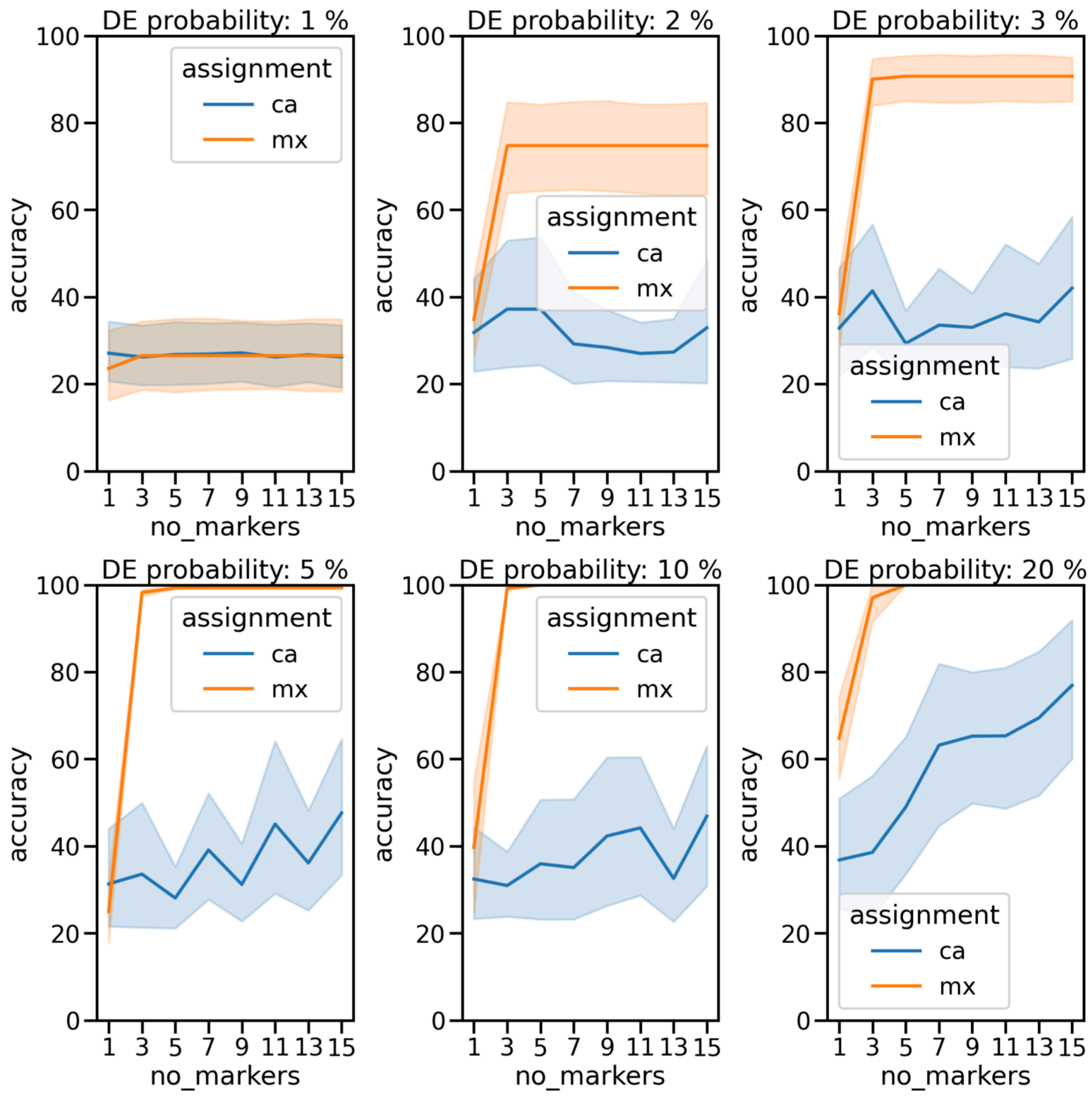


**Supplementary Figure 3:** The same approach as that described in Supplementary Figure 1 was used, but 15 marker genes were chosen for all DE probabilities by selecting genes with the highest log-fold change among those that were in the top tenth percentile of mean expression.

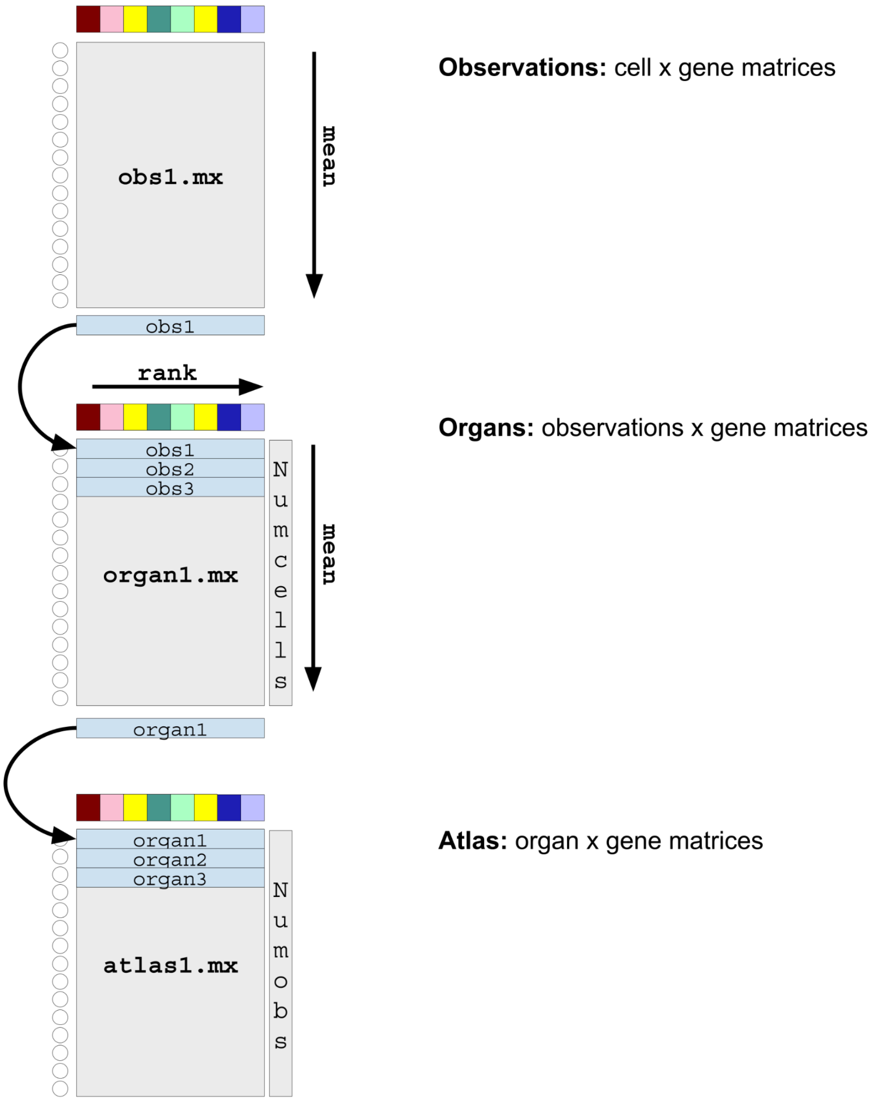


**Supplementary Figure 4:** Summarizing atlas content. Gene counts are averaged across cells within individual cell-by-gene matrices to create "organ" level summaries of count matrices. Averaged counts are ranked across genes and then subsequently averaged to create whole-atlas-level summaries of count matrices.

Zhang, Allen W., Ciara O'Flanagan, Elizabeth A. Chavez, Jamie L. P. Lim, Nicholas Ceglia, Andrew McPherson, Matt Wiens, et al. 2019. "Probabilistic Cell-Type Assignment of Single-Cell RNA-Seq for Tumor Microenvironment Profiling." *Nature Methods* 16 (10): 1007–15.
